# Powerful read processing with *matchbox*

**DOI:** 10.1101/2025.11.09.685711

**Authors:** Jakob Schuster, Kathleen Zeglinski, Lucinda Xiao, Olivia Voulgaris, Sarahi Mendoza Rivera, Stephin J. Vervoort, Matthew E. Ritchie, Quentin Gouil, Michael B. Clark

**Affiliations:** The Walter and Eliza Hall Institute of Meidcal Research, Parkville, 3052, Victoria, Australia; Department of Medical Biology, The University of Melbourne, Parkville, 3052, Victoria, Australia; Department of Anatomy and Physiology, The University of Melbourne, Parkville, 3010, Victoria, Australia; Olivia Newton-John Cancer Research Institute, Heidelberg, 3084, Victoria, Australia; School of Cancer Medicine, La Trobe University, Bundoora, 3086, Victoria, Australia

## Abstract

The wide variety of protocols and applications for DNA and RNA sequencing makes flexible tools for read processing an important step in sequence analysis. Beyond trimming and demultiplexing, custom read-level processing is commonly required for data exploration, QC and analysis. Existing tools are often task-specific and don’t generalise to new bioinformatic problems. Thus, there is a need for a tool flexible enough to handle the full variety of read processing tasks, and fast and scalable enough to retain high performance on growing sequencing datasets. We introduce *matchbox*, a read processor that enables fluent manipulation and analysis of FASTA/FASTQ/SAM/BAM files. With a lightweight scripting language designed around error-tolerant pattern-matching, users can write their own *matchbox* scripts to tackle a wide variety of bioinformatic problems, and incorporate them into existing pipelines and workflows. We demonstrate *matchbox*’s versatility in a number of contexts: demultiplexing long-read scRNA-seq data with 10X or SPLiT-seq barcodes; restranding RNA-seq reads; assessing CRISPR editing efficiency; and haplotyping macrosatellite repeat regions. *matchbox* achieves a computational performance comparable to existing tools, while addressing a broader range of bioinformatic needs, representing a new state-of-the-art in sequence processing. *matchbox* is implemented in *Rust* and available open-source at https://github.com/jakob-schuster/matchbox.

## Introduction

Read processing is an important step after basecalling of sequencing reads and can involve read QC and filtering; exploring read structures and read-level analysis; and preparing reads for downstream analysis. Some read processing steps are common across sequencing methods and platforms, such as trimming off synthetic primers or demultiplexing based on sample or cell barcodes. However, even these routine tasks become complex in certain experimental contexts, such as when dealing with combinatorial barcoding schemes (6). Read-level analysis tasks can also be highly specialised, even experiment-specific; for example, when a novel combination of protocols or techniques is used (6), or when read structures are unique to the experiment (7).

Several recent tools have been developed in response to this growing need for flexibility. *flexiplex* uses command-line syntax to specify a configuration of primer, barcode and universal molecular identifier (UMI) regions (8). While the tool is ostensibly for demultiplexing reads, it can also perform error-tolerant sequence searches, as well as various custom trimming and demultiplexing-adjacent tasks. However, although there is great flexibility in the read structures it can parse, *flexiplex* only supports a few predefined operations and many complex tasks require multiple consecutive applications of *flexiplex*, incurring a computational cost.

Another recent approach, *splitcode*, utilises a configuration file, which affords it more programmability (9). The user can specify each fixed sequence or barcode to search for, the order in which to perform the searches and the edit distance of each search. Then, they can describe a number of operations to perform (e.g. after the primer sequence, extract the following 8 base pairs and place them in the description line of the read). With this programmable behaviour, *splitcode* is more flexible than *flexiplex*, however the configuration format still creates limitations to its generality. In addition, the implementation details of its search algorithm mean *splitcode* cannot search for long sequences or tolerate high error rates, rendering it incompatible with many long-read applications.

Existing approaches are thus limited by their interfaces. *flexiplex*’s command-line syntax is simple and readable, but does not afford the expressiveness to perform complex read manipulation. *splitcode* allows for deeper specification, but it sacrifices readability in its configuration format, which consists of a tab-delimited table. Many bioinformatic ideas which are difficult to implement using existing tools can be easily expressed in terms of conditional pattern-matching (e.g. first check if a read has a particular structure; if so, trim it and output it; otherwise, check if it has another particular structure) and function application (e.g. reverse-complement a sequence; annotate a read with some metadata). Thus, a general-purpose tool supporting this kind of structured expression of bioinformatic intent would be more flexible and its configuration would be more readable compared to existing tools.

By introducing purpose-fit, domain-specific programming languages, *snakemake* and *Nextflow* have improved the expeience of bioinformatic workflow development (10) (11). In the same sense, a tool which approaches read processing as a language design problem could empower bioinformaticians to handle complex read-level analysis tasks. By using a simple scripting language as its interface, a read processing tool could provide extensive flexibility without sacrificing readability.

To be applicable to massive sequencing datasets, a read processor must also be implemented with speed in mind. *Rust* is a modern systems programming language with excellent performance and a growing variety of bioinformatics packages (12). It has become increasingly popular for implementing compute-intensive bioinformatics tools (13) (14), and its modern design features make it well suited to the implementation of a domain-specific language compared to older languages such as *C* or *C++*.

Here we present *matchbox*, a new paradigm for flexible and efficient read processing based on a domain-specific scripting language. We demonstrate *matchbox*’s versatility, applying it to routine tasks such as error-tolerant sequence searching and cell demultiplexing, as well as tailored analyses of CRISPR prime-editing libraries and macrosatellite repeats in genomic data. We also benchmark the performance of *matchbox*, finding that the tool achieves a high throughput equivalent to existing purpose-built tools, while providing greater flexibility.

## Materials and methods

### A. Implementation

*matchbox* is implemented in *Rust. matchbox* execution can be divided into compile time, which involves taking in the user’s script, and runtime, which involves processing the reads based on the instructions given by the user.

#### A.1 *Compilation of* matchbox *scripts*

When the user starts *matchbox*, their *matchbox* script is parsed into a surface-level abstract syntax tree (AST), which closely reflects the written syntax of the language. The surface AST is elaborated to a core AST, which can be thought of as a concrete series of instructions interpreted at runtime, evaluating for each read a set of output effects (e.g. writing a trimmed read to a file, printing to standard output, incrementing a global count). During the process of elaboration, bidirectional type-checking is performed. Type-checking ensures that functions have been given the right type of arguments; for example, the `len` function must take `Str` input. Bidirectional type-checking means checking the types of terms with known types, while inferring the types of terms with implicit or unknown types. This means *matchbox* users do not need to annotate their types, or indeed be familiar with types at all, despite benefiting from their guarantees.

Then, *matchbox* identifies terms that do not depend on the value of an input read, evaluates them, and stores their values so they do not need to be evaluated again. For example, the term `csv(‘large_csv_file.csv’)` is evaluated at compile time, to avoid reopening and parsing the CSV every time a read is processed. This caching ensures that the minimum amount of computation is done at runtime when processing reads.

#### A.2 Multi-threaded execution

During runtime, processing in *matchbox* is parallelised at the read-level. A chunk of the input file is read into memory and the work is executed in parallel across a thread pool using the *rayon* data parallelism library (15). Then, the results are aggregated and any output is produced before the next chunk of reads is processed. The `-threads` parameter determines the size of the thread pool.

#### A.3 Error-tolerant pattern matching

Aligning sequences while allowing for edit distance is the most computationally intensive part of *matchbox*. All pattern matching uses Levenshtein distance, and is built on the *Rust-bio* implementation of Myers’ bit-parallel approximate string matching algorithm (16) (12). Since the algorithm does not involve precomputation, as *splitcode*’s or *ugrep*’s algorithm does, *matchbox* can handle high error rates without incurring computation and memory costs at startup (9).

When matching a pattern consisting of several regions, *matchbox* aligns the regions one by one, each time constraining its next search to the range allowed by previous results. For example, to match a single read against the pattern `[_ prim1 _ prim2 _]`, *matchbox* will first locate the `prim1` sequence, before searching for `prim2` only in the part of the read that comes after where `prim1` was found. In the case that one pattern binds to multiple places in the read, e.g. the read has the structure `[_ prim1 _ prim2 _ prim1 _ prim2 _]`, the `-match-mode` parameter determines whether to consider all matches, or only the one with the lowest cumulative Levenshtein distance.

### B. Benchmarking

#### B.1 Datasets

To demonstrate *matchbox*’s functionality, various existing datasets were used, summarised in Supplementary Table 1 (1–5, 17–20).

#### B.2 Software

A full list of software used is given in Supplementary Table 2 (1, 3, 8–10, 12, 15, 17, 18, 21–31). All analysis scripts are also available on GitHub at https://github.com/jakob-schuster/matchbox-analysis-scripts.

#### B.3. Performance benchmarking

All tools were given 32 cores and 32 GB of RAM. Where tools included a parameter for multithreading, they were given 32 threads. *hyperfine* was used to benchmark the tools, running each one 3 times and averaging across the runs. For all tasks, samples of 10M reads were used, hence throughput in reads/sec was calculated as (10M / total time). Benchmarks were run on a high performance computer using Intel(R) Xeon(R) E5-2690 v4 @ 2.60GHz (Broadwell) CPUs. SSDs used for reading and writing input files were VAST high performance all-flash NVMe storage.

## Results

### C. Syntax and usage

*matchbox* can be run from the command line. The user provides reads to be processed, as well as a *matchbox* script describing what they want to do (Figure 1A). Some general settings are available as command-line parameters, including the global error rate applied to all search operations. A *matchbox* script is a series of statements, executed from top to bottom for each input read (Figure 1B). Many typical programming language features are present in *matchbox*: variables can be assigned for re-use, conditional `if` statements can be used to execute code when certain conditions are met, and functions can be used to manipulate data. Some functions, marked with an exclamation!, produce output – for example, the `stdout!()` function prints a value directly to the standard output, the `out!()` function writes a value to a file, and the `mean!()` function allows the user to compute the mean of a numerical value across all reads, such as read length.

**Fig. 1.**
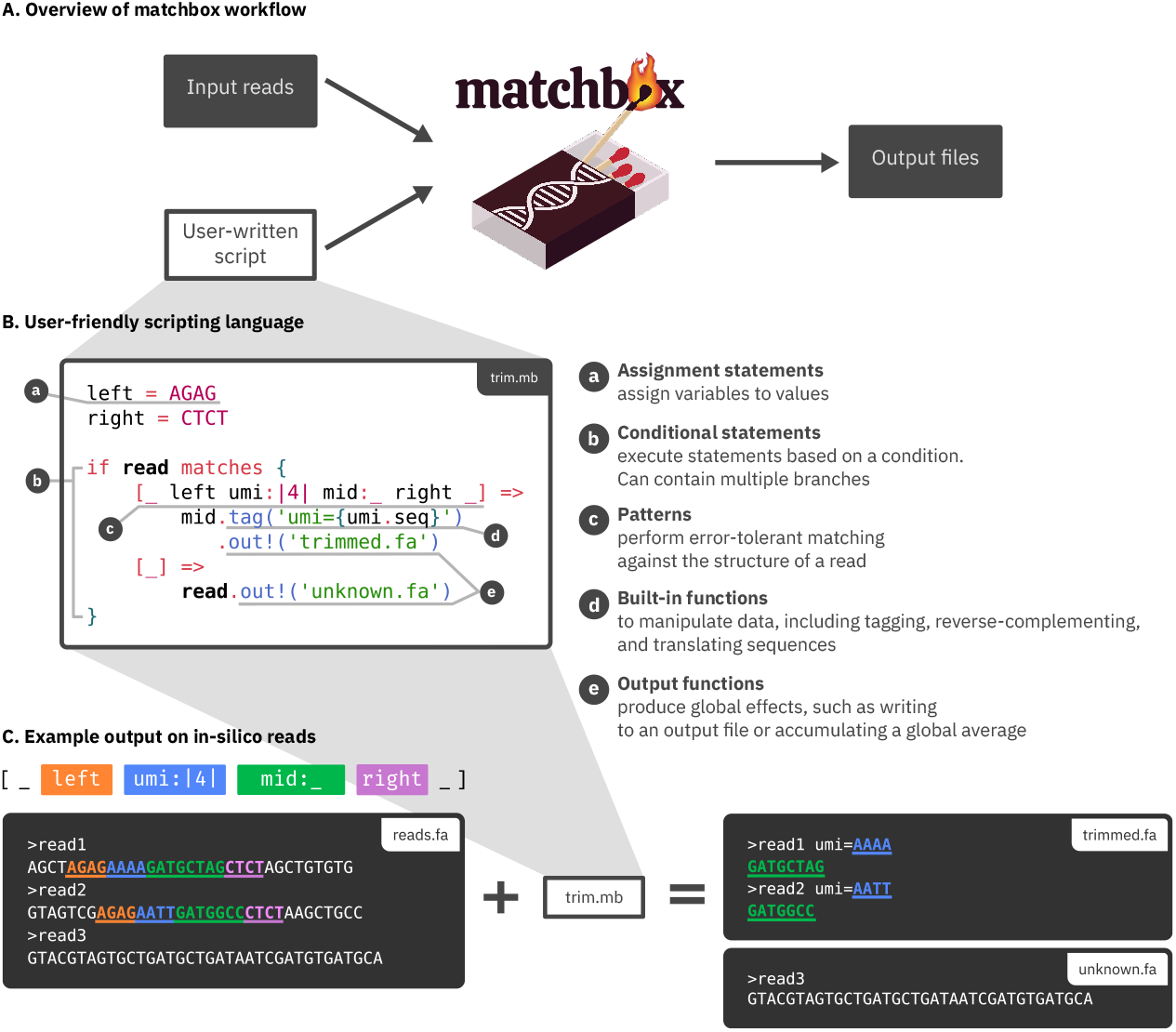
Overview of *matchbox* usage and syntax. (A) *matchbox* is run on the command line, taking as input a read file in FASTA/FASTQ/BAM format and a script describing the processing steps to perform on each read. Depending on the user’s script, *matchbox* produces various outputs, including summary statistics, information extracted from each read, or modified read files in FASTA/FASTQ/BAM format. (B) An example script called `trim.mb` which trims off a left-flanking primer `AGAG` and a right-flanking primer `CTCT` . A 4-bp UMI is also extracted following the left-flanking primer, and the trimmed read is tagged with the UMI in its description line before being printed to the output file `trimmed.fa.` Reads which do not contain the two primer sequences are sent to a file `unknown.fa`, so that they can be investigated further. (C) The result of running the `trim.mb` script on three FASTA reads. Colour coded schematic of the pattern to be matched and corresponding nucleotides are displayed. `read1` and `read2` contain the expected primers and are included in the output `trimmed.fa`, tagged with their respective UMI sequences. Since `read3` did not contain the primers, it is included in `unknown.fa` .

The core feature of *matchbox* is pattern-matching of reads. Using the keyword `matches`, the user can write out a series of patterns to test each read against (Figure 1B). Only the first successfully matching pattern executes the statements in its branch. A pattern is a schematic describing the structure of a read from left to right, comprised of a series of regions (Figure 1B, C). Variable regions marked with an under-score _ represent any number of any nucleotides; they will match anything. Fixed-length regions marked with vertical bars `|n|` represent an exact number of nucleotides. When a known sequence or a variable name is included in a pattern, the sequence is searched error-tolerantly in the read; if the sequence cannot be found, the read does not match the pattern. Naming a region using a colon : allows the sequence to be extracted and treated as a slice. Patterns can also include parameters using the keyword `for .. in`, whereby a variable is bound to one option from a list (e.g, a single barcode from a barcode list). By combining these basic regions, a wide variety of read structures can be described. For a more in-depth explanation of the *matchbox* scripting language, the documentation site provides a comprehensive reference with examples.

### D. Basic examples

*matchbox* can be used as an efficient error-tolerant grep, demonstrated here by searching for a template-switching oligo (TSO) primer in an Oxford Nanopore (ONT) PCR-cDNA dataset (Figure 2A) (1). When pattern-matching, *matchbox* tolerates error as a proportion of each searched sequence’s length, given globally by the `-error-rate` parameter. For example, an error rate of 0.2 applied to the 26-bp TSO would allow a maximum edit distance of 5 (rounded to the nearest integer). Since this dataset contains reads representing first- and second-strand cDNA, we expect to find this TSO primer on roughly 50% of reads, with the remaining reads containing the reverse-complement of the primer sequence. As *matchbox*’s error rate is increased to 0.2, the proportion of grepped reads approaches the expected 50% (Figure 2B). However, when the error rate is in-creased above 0.3, the proportion of reads quickly rises above the expected proportion. This indicates that, while some error tolerance picks up valuable signal, too much will detect noise. The appropriate error rate setting for any task will thus depend on the bioinformatic and biological context.

**Fig. 2.**
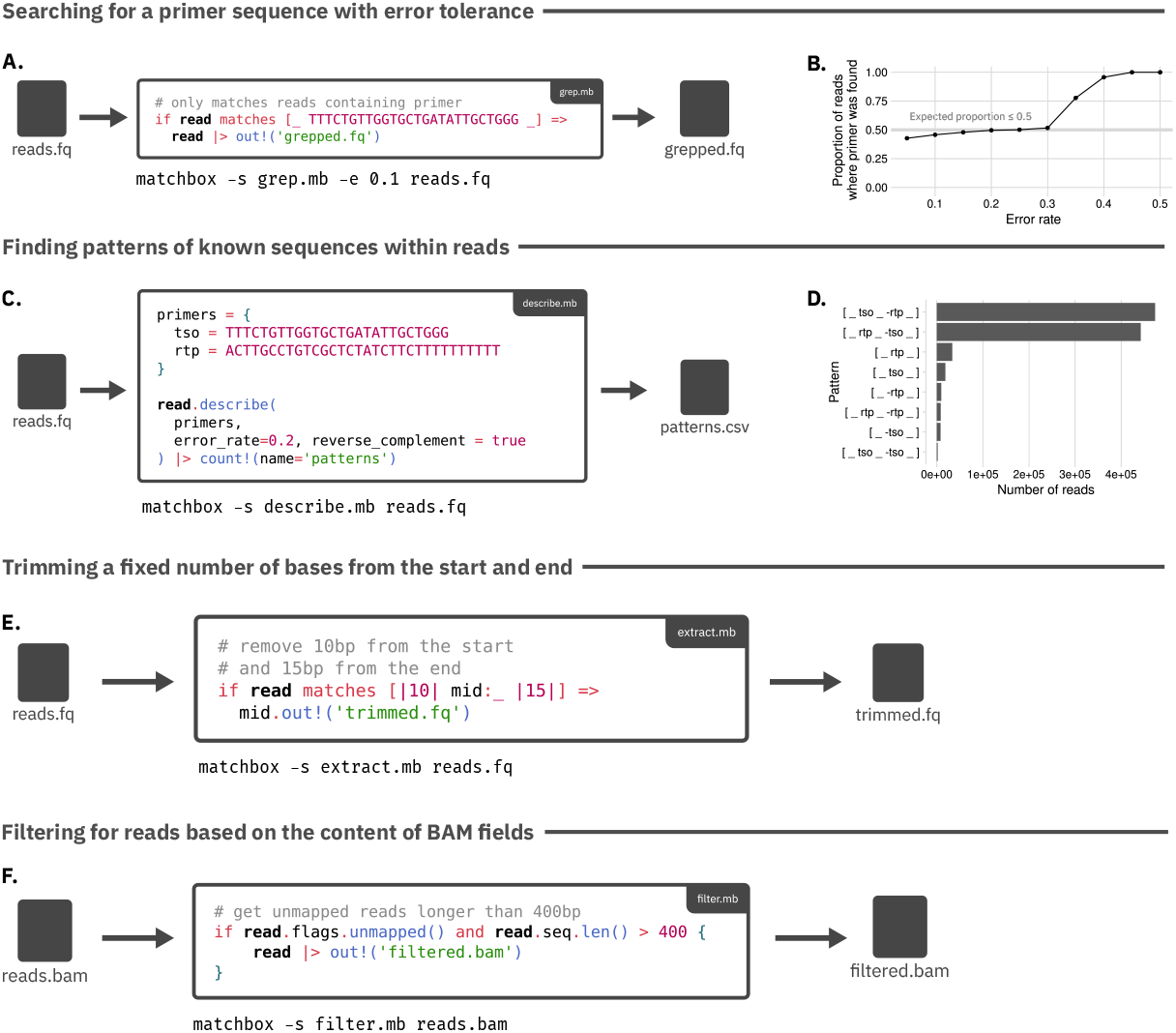
Examples of common *matchbox* use cases, applicable to many kinds of sequencing data analysis. (A) *matchbox* script to filter reads containing a TSO primer sequence into `grepped.fq` . Error rate is configured via the `-e` command line option. (B) Impact of changing *matchbox* ‘s maximum error rate on proportion of matching reads, demonstrated on an ONT PCR-cDNA sequencing dataset (1). Since the dataset contains reads representing first- and second-strand cDNA, the TSO sequence is expected to be found on approximately 50% of reads. (C) Script to find the arrangements of TSO and RTP sequences in a dataset. The optional argument `reverse_complement` indicates that each sequence should also be searched for in the reverse-complementary direction. (D) The most frequent patterns found in the dataset, with rare patterns (*<*1000 reads) removed during visualisation. The two most common patterns are the expected forward and reverse read structures. (E) *matchbox* script for trimming off the first 10 bp and last 15 bp from each read. (F) *matchbox* script to filter for unmapped reads *>*400 bp in length.

*matchbox* can also be used to discover the arrangements of known sequences in reads, a useful feature when exploring an unfamiliar dataset or diagnosing artefact-related issues in an experiment (Figure 2C). Known sequences are supplied to the `describe()` function, which searches for each primer and describes the read in terms of the sequences provided. Optional arguments to the function allow error rate to be tuned and allow for the reverse-complement of each sequence to also be searched. Run on the same PCR-cDNA dataset, the most frequent patterns `[_ tso _ -rtp _]` (RTP: reverse transcription primer) and `[_ rtp _ -tso _]` represent the expected forward and reverse read structures (Figure 2D). Other structures are also reported, including reads where only one primer could be found, as well as TSO arte-facts `[_ tso _ -tso _]` and RTP artefacts `[_ rtp _-rtp _]`, known structures resulting from RT-PCR errors (1).

*matchbox* can be used to extract or remove fixed numbers of bases, here used to trim 10 bp from the start and 15 bp from the end of a read (Figure 2E). If a read is less than 25 bp in length, the pattern will not match, and the read will not be included in the ‘trimmed.fq’ output.

Additionally, *matchbox* can process SAM/BAM files, and allows for reading and writing SAM flags and optional tags.

Here, unmapped reads longer than 400 bp are extracted (Figure 2F).

We will now outline several diverse read processing scenarios, comparing *matchbox* to existing tools and highlighting its more advanced features.

### E. Demultiplexing single-cell data

Sequencing data is often multiplexed via the addition of synthetic barcode sequences during library preparation. These barcodes are used to distinguish reads originating from separate samples or, in the case of single-cell sequencing, from separate cells. This means a crucial step in analysis is demultiplexing: locating barcode sequences within reads and assigning them to the correct cells and/or samples (32).

In the 10X single-cell barcoding approach, each read receives a single 16-bp cell barcode (Figure 3A) (32). Since searching error-tolerantly for all 3.3 million 10X barcode sequences in each read would be computationally impractical, the first step in demultiplexing long-read data is determining the subset of 10X barcodes present in the data that represent real cells; this step is known as barcode discovery (3). A simple *matchbox* discovery script can be devised, extracting 16-bp barcodes following a primer sequence and reporting the frequencies of each barcode’s occurrence (Figure 3B). This list can then be filtered using an existing tool *flexiplex-filter* or any heuristic to remove low-frequency barcodes (which represent sequencing errors or empty droplets), to produce a final filtered list (8). Testing on multiple long-read scRNA-seq datasets, which also contained matched short-read scRNA-seq to act as positive controls for barcode identification, the *matchbox* script produced high quality barcode lists with apparent false discovery rates of 0%, 3.89% and 1.67% across the three datasets (Figure 3C). The number of cells detected was highly congruent with existing tools, despite the script being extremely simple and only a few lines in length.

**Fig. 3.**
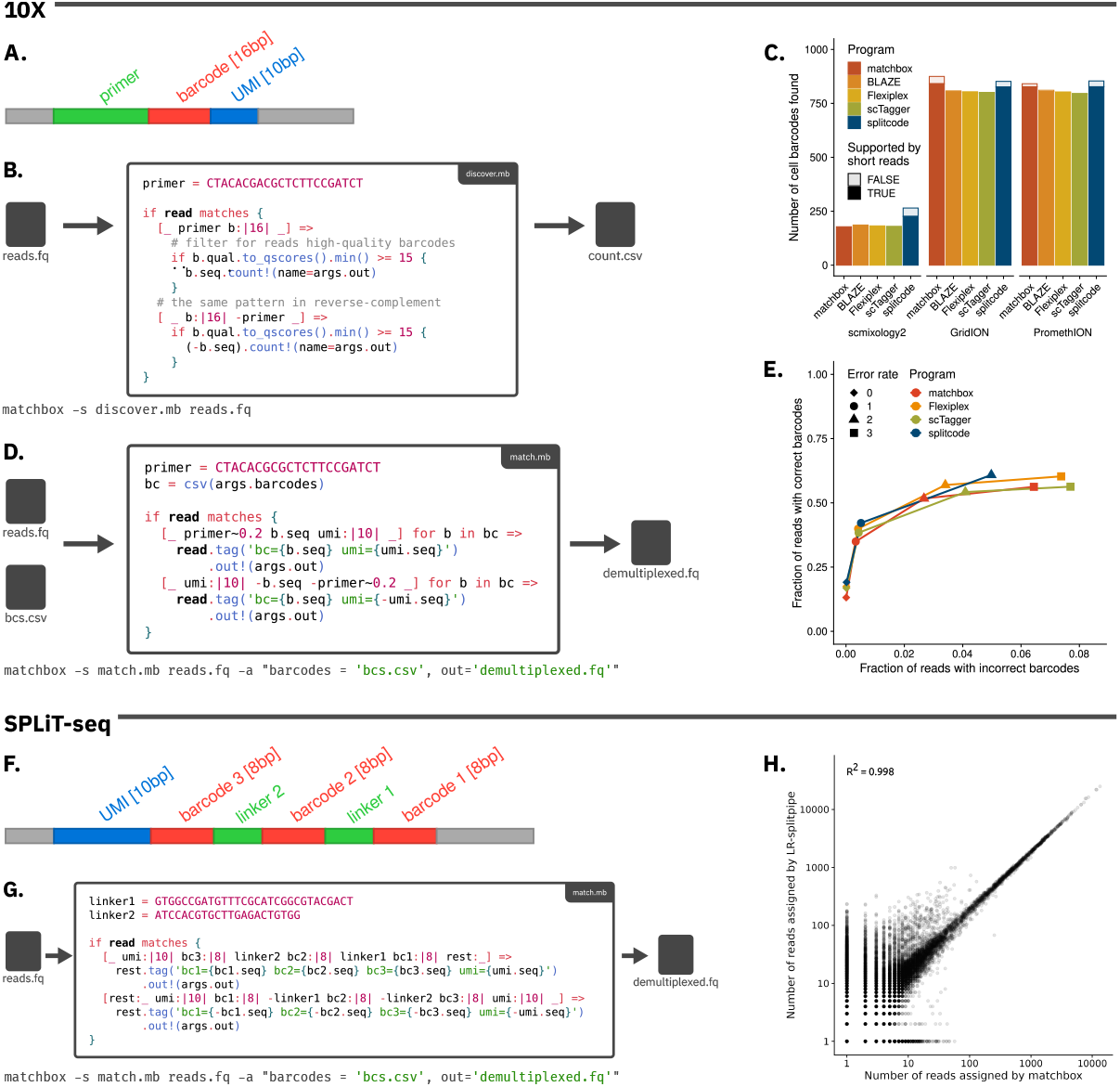
Application of *matchbox* to demultiplexing cell barcodes. (A) Schematic of reads barcoded with 10X method. (B) *matchbox* script for 10X barcode discovery. For each read, the primer is located and the following 16 bp are extracted. Occurences of each barcode are counted and reported in `bcs.csv` . (C) Comparison of filtered barcode lists generated by barcode discovery tools on the scmixology2 dataset (2), as well as a hiPSC dataset sequenced on the ONT GridION and PromethION (3). For both datasets, barcode lists derived from paired short-read data were used to assess how accurately tools performed discovery. (D) *matchbox* script for demultiplexing 10x reads using a list of known barcodes. Primers are searched using a fixed error rate of `0.2`, and barcode error rate is varied with the command line parameter `-e` . Barcodes and 10-bp UMIs are extracted from each read and placed as tags in the description line of the FASTQ output. (E) Precision and recall of barcode matching tools on a simulated dataset where the true barcode for each read was known. When the edit distance was set higher than 2, *splitcode* could not be tested as it required more memory than we could provide (*>*256 GB). (F) Schematic of reads barcoded with SPLiT-seq method. (G) *matchbox* script for demultiplexing SPLiT-seq barcodes. Barcodes are naively extracted from reads without comparison to a reference. (H) Number of reads assigned to each barcode by *matchbox* and existing tool *LR-splitpipe*.

Once a list of barcodes has been discovered, the list can be used to error-tolerantly assign each read to its barcode. Again, the *matchbox* approach is very simple (Figure 3D), yet on a simulated dataset where the true barcode assignment of each read is known, it performed similarly to existing tools (Figure 3E).

The SPLiT-seq (also known as PARSE) single-cell barcoding approach differs from the 10X method, resulting in a complex 3-barcode scheme that must be computationally demultiplexed (Figure 3F) (6). In particular, for long-read SPLiT-seq, where barcodes can only be located by their adjacency to linkers and not based on absolute position in the read, the number of tools that support the schema are limited; 10X-specialised tools such as *scTagger* are unable to parse the barcoding scheme (18). A *matchbox* script can easily be written to handle this format (Figure 3G). In this naive approach, the barcodes were extracted directly from the reads and not checked against the barcode reference file. Compared to specialised tool *LR-splitpipe* (18), the *matchbox* script assigned each read the exact same barcode as *LR-splitpipe* 86% of the time, and the read counts per barcode between *LR-splitpipe* and the *matchbox* script were well-correlated with an *R*^2^ of 0.998 (Figure 3H). In summary, *matchbox* scripts offer flexible, simple demultiplexing for complex barcoding schemes, with similar performance to specialised demultiplexing tools.

### F. Restranding PCR-cDNA data

RT-PCR results in first- and second-strand cDNA copies of each RNA transcript, obscuring its original strandedness. However, since the reverse transcription primer (RTP) and template-switching oligo (TSO) are annealed prior to PCR, the reads representing first- and second-strand cDNA can be distinguished by the presence of forward or reverse primer structures (Figure 4A). The first-strand reads can then be reverse-complemented so that all of the transcripts are oriented in their original direction, a process known as ‘restranding’. TSO and RTP artefacts also result in differently structured reads, which can be filtered out during such preprocessing.

**Fig. 4.**
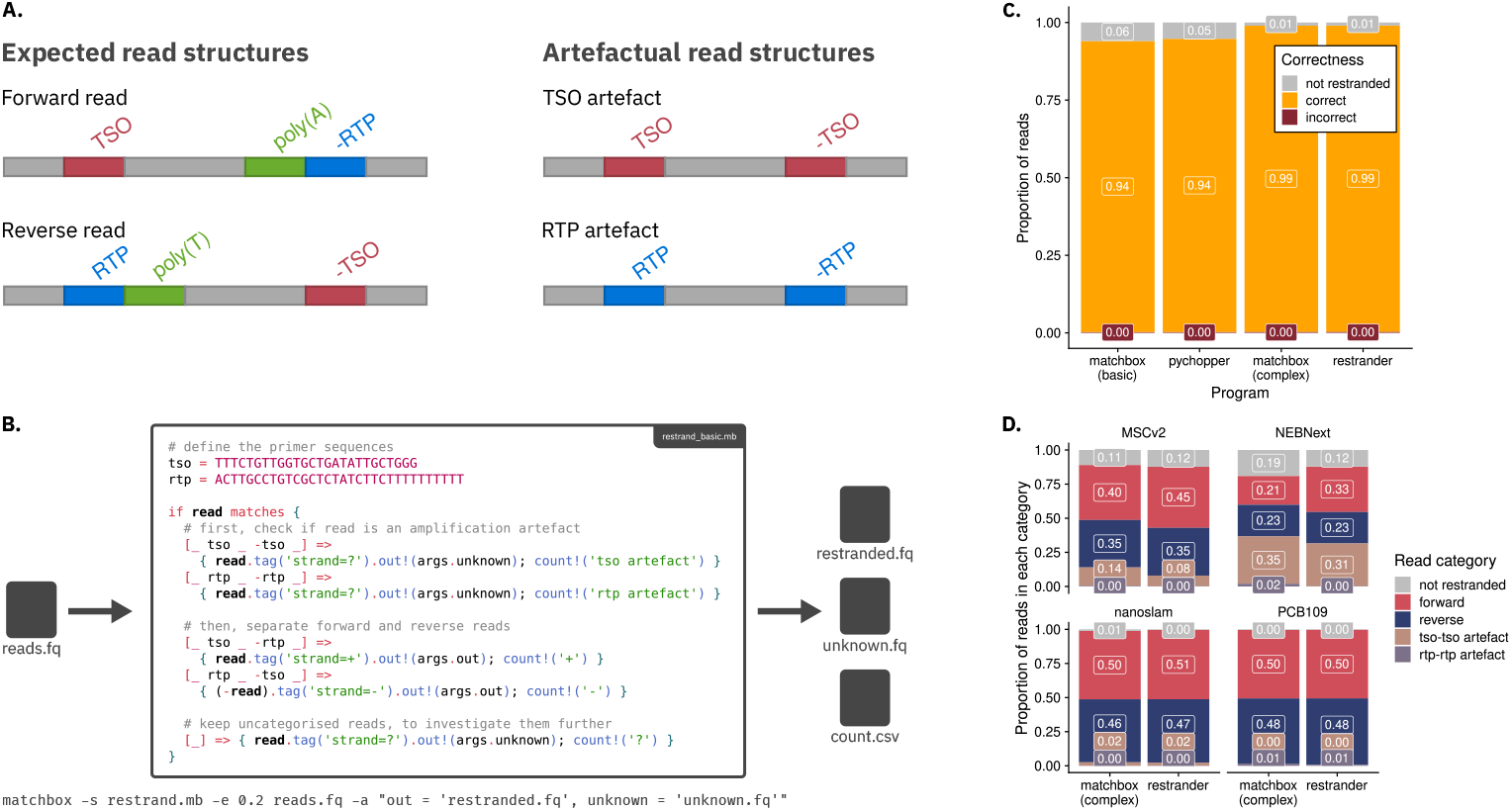
Application of *matchbox* to restranding sequencing reads. (A) Schematic of read structures resulting from ONT PCR-cDNA sequencing. RTP - reverse transcription primer; TSO - template-switching oligo; poly(A) - poly(A) tail. “-” symbols indicate reverse-complemented sequences. (B) The *matchbox* script to perform restranding. Each read is matched against patterns representing forward and reverse primer structures, as well as artefactual structures, and sent to `restranded.fq` and `unknown.fq` accordingly. Reads of each category are also quantified, with the results collected in a CSV file. When searching for primers, an error rate of `0.2` is permitted. (C) Comparison of *matchbox, Restrander* and *pychopper* on the subset of reads mapped to sequins (synthetic transcripts whose strandedness is known). Tools either inferred the strand of a read correctly, incorrectly, or opted not to make a judgement. (D) Demonstration of the complex *matchbox* script on various libraries with different primer sequences, including mouse muscle stem cells prepared with 10X Genomics Chromium 3’ kit (2), rat single B cells prepared with NEBNext Low input/single cell kit (4), mouse muscle stem cells prepared with ONT PCB111 (1) and the lung adenocarcinoma cell lines prepared with ONT PCB109 (5).

Multiple dedicated tools exist for restranding. *Pychopper*, developed by ONT, is a *python*-based tool which detects primers and classifies reads as either forward (second-strand) or reverse (first-strand) (22). Using a simple *matchbox* script, we can perform restranding in the same way, using concise pattern-matching syntax to separate forward, reverse, and artefactual reads and process each read type accordingly (Figure 4B). A more recent tool, *Restrander*, applies a more involved algorithm than *pychopper*, first attempting to restrand reads based on the presence of poly(A) or poly(T) tails near the start and end of reads before searching for primers (1). A more complex *matchbox* script was also written to mimic this behaviour (Supplementary Methods 3.2).

To assess the performance of the *matchbox* scripts against existing tools, a Nanopore longread PCR cDNA-seq dataset was used which included sequins (synthetic transcripts with known strandedness) (5). The reads were aligned to the sequin reference using *minimap2*, and the `ts` flag (used by *minimap2* to indicate a read had to be reverse-complemented during alignment) was used as a read-level ground truth. On the subset of sequin-aligned reads, all methods correctly restranded over 90% of reads (Figure 4C). The simple *matchbox* approach performed similarly to *pychopper*; this was unsurprising since both approaches base their classifications on a simple error-tolerant primer search. The more complex *matchbox* script achieved superior restranding, close to the performance of *Restrander*. We applied the complex *matchbox* script to a variety of protocols with different primer sequences (Figure 4D).

As illustrated by this example, 17 lines of a *matchbox* script can achieve the same result as 1400 lines of *C++* in *Restrander*. Additionally, the basic bioinformatic logic of restranding is easy to understand by reading the script, and different approaches to the problem (the simple vs complex approach) are quick to develop and compare, a kind of exploration that would be time consuming to implement when developing conventional tools. Modifications to *matchbox* scripts are also much less costly; tweaking the script so that primers are trimmed off during restranding would take less than a minute (Supplementary Methods 3.3), whereas the same change may take several hours or days to implement correctly into a conventional tool. This highlights the power, simplicity and generalisability of *matchbox*.

### G. Analysing self-targeting CRISPR screens

Many experimental techniques require tailored analysis of complex oligonucleotides with known structures; one example is CRISPR prime-editing self-targeting screens. Prime editing requires two components: a guide RNA (pegRNA), which targets a genomic location and encodes the intended edit, and a Cas9 protein that performs the editing (7). Since editing rates are often highly variable between individual guides, assessing the editing efficiency of each one is essential. Editing efficiency can be assessed via a self-targeting screen, whereby pegRNAs contain both a targeting sequence and the target sequence to be edited, enabling Cas9 to edit the pegRNA in addition to the genomic target. Sequencing of pegRNAs therefore quantifies their editing efficiencies, as self-targeting efficiencies have been shown to match endogenous genomic editing efficiencies (19).

Cirincione and Simpson *et al*. (2024) performed a self-targeting CRISPR screen to assess the editing efficiency of each pegRNA guide over time (7-28 days), between cell lines (PEmax vs PEmaxKO) and across each substitution type (G*>*A, G*>*C, G*>*T) (Figure 5A) (19). Each guide had a unique barcode, as well as a target sequence with an expected edit (Figure 5B). For each guide, editing efficiency was calculated as the proportion of reads containing the edited target sequence. Since the read processing required was dependent on the structure of the pegRNA, existing tools were insufficient and the authors resorted to a custom *python* script. For this type of tailored oligo-specific analysis, *matchbox* is highly applicable.

**Fig. 5.**
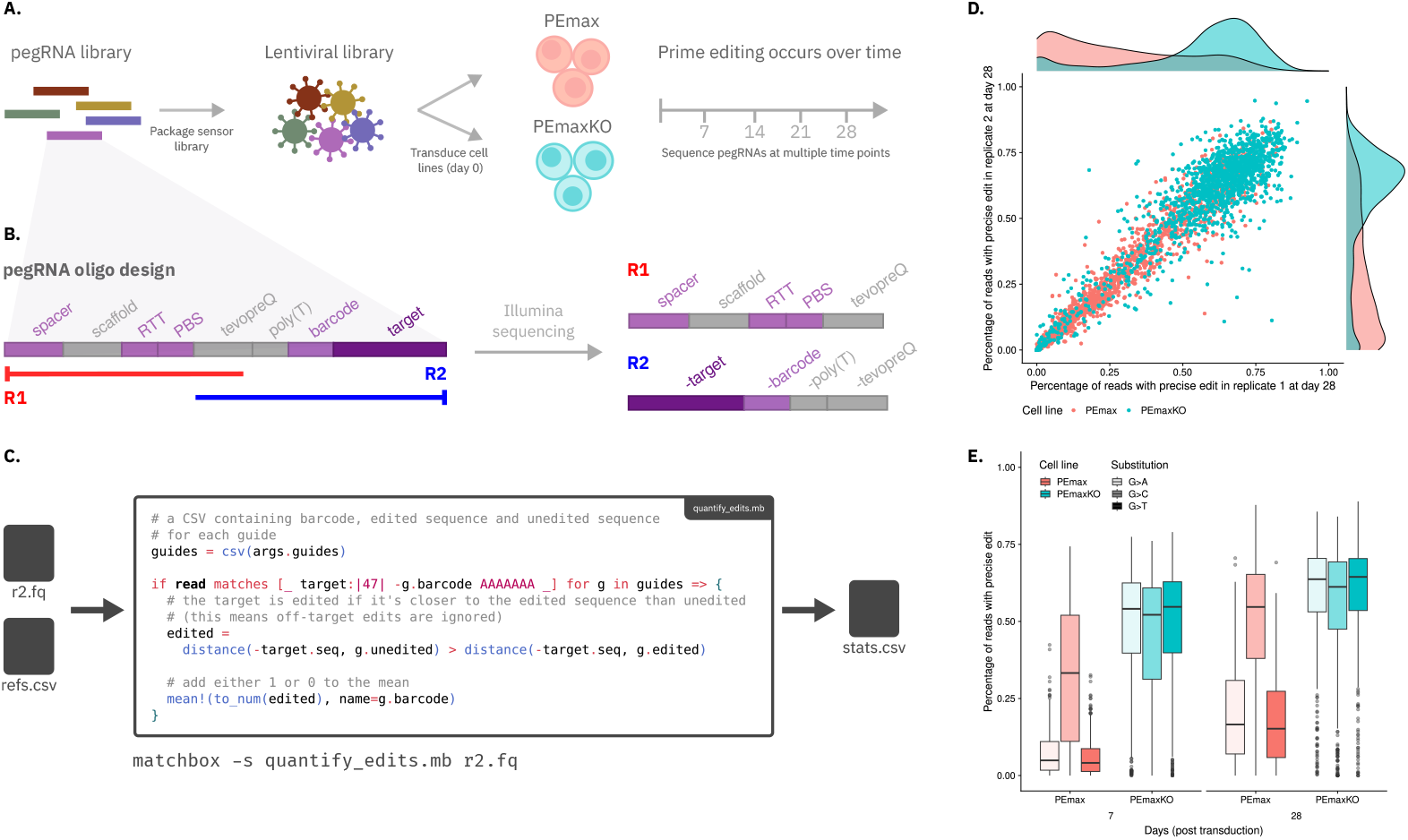
Application of *matchbox* to analysis of a self-targeting CRISPR screen. (A) Workflow of a self-targeting CRISPR screen. A library of prime editing guide RNAs (pegRNAs) were packaged into a lentiviral vector and transduced into two cell lines. Sequencing to measure editing efficiency was then performed at weekly intervals. (B) pegRNAs include regions of fixed (gray) and variable (purple) sequence. The spacer, reverse transcriptase template (RTT) and primer binding site (PBS) regions specify the targeted sequence and the intended edit. Each guide in the library is identified by a unique barcode. The target region matches the genomic target sequence and is edited, allowing the editing efficiency to be determined by sequencing the oligos. R1 and R2 represent Illumina sequencing reads 1 and 2 and the portion of the pegRNA they cover. R2 includes both the target and the barcode, all the information required to assess editing efficiency. (C) The *matchbox* script for assessing editing efficiency. For each read, the barcode is located (in reverse-complementary direction, since this is R2). Then, it is determined whether the target region is more similar to the edited or the unedited sequence. Either a 1 or a 0 is appended to a global average for that barcode, and a CSV is produced recording the average edit frequency of each barcode. (D) Scatter plot showing the proportion of edited reads for each guide at day 28, comparing across both replicates and the two cell lines PEmax and PEmaxKO in Cirincione and Simpson *et al*’s ‘+5 G*>*H’ library. (E) Proportion of edited reads for each guide over time, averaged across replicates, grouped by substitution type and compared across cell types PEmax and PEmaxKO.

Using a short and simple *matchbox* script we replicated the results of Cirincione and Simpson *et al*. (2024), demonstrating editing efficiency was higher in PEmaxKO cells (Figure 5C, D). Additionally, editing took place much more quickly in PEmaxKO cells, with a majority of edits having occurred by 7 days post transduction (Figure 5E). While PE-max cells exhibited very low editing efficiency for G*>*A and G*>*T substitutions, PEmaxKO cells displayed more consistent editing across substitution types (Figure 5E).

The *matchbox* script assigns each read to its barcode, then classifies it as either edited or unedited (Figure 5C). The barcode of a read is matched to one guide in the reference CSV file, and the read’s 47-bp target sequence is extracted. If the extracted target sequence is closer to the matching guide’s edited sequence than to its unedited sequence (in terms of Levenshtein edit distance), the read is considered edited. Alternative approaches to calculating edit distance include filtering out reads with off-target edits, or classifiying them in their own read category and quantifying the prevalence of each off-target edit quantified for analysis. These alternative implementations were also written up as *matchbox* scripts (Supplementary Methods 4.1). Importantly, the specific editing calculation used for the analysis is not buried in a several-hundred-line *python* script, but exposed in a concise 11-line *matchbox* script. By reducing the script to only its bioinformatic concept, *matchbox* makes the expression of complex bioinformatic ideas in code simpler.

### H. Analysing a macrosatellite repeat region

Haplotyping of human genomic data in the context of genetic disorders can require tailored analyses at particular loci. One example is the D4Z4 macrosatellite array on chromosome 4, the genetic and epigenetic dysregulation of which is associated with facioscapulohumeral muscular dystrophy (FSHD) (33). The D4Z4 array has a known genetic structure, consisting of a constant proximal flanking sequence, followed by 1 to *>*100 3.3kb tandem repeats, followed by one of two distal sequences (A or B), which define the two 4q haplotypes, 4qA and 4qB. A full-length *DUX4* gene is present at the end of the array, and is silenced in healthy muscle cells. A 4qA array with 1-10 repeat units (RU) is associated with pathogenic *DUX4* expression, and causes the most common type of FSHD, FSHD1 (95% of cases) (33). A homologous qA-type D4Z4 array is also present on chromosome 10q, which can be distinguished from the 4q array by slight sequence differences including a 10q-specific BlnI site, and a base substitution in the qA sequence that renders it non-pathogenic. Genetic testing for FSHD therefore requires accurate ‘matching’ of multiple features, for chromosome 4q/10q assignment, A/B haplotyping, and D4Z4 repeat number counting.

Recently, long-read Nanopore sequencing was applied to the D4Z4 array, and the *D4Z4End2End* pipeline developed for genetic and epigenetic analysis of FSHD (23). To haplotype the D4Z4 locus, reads originating from each allele can be identified from reads that span the whole array (including the proximal and distal flanking sequences). Then, in spanning reads, D4Z4 repeats are counted to identify pathogenic alleles. *D4Z4End2End* involves a custom *python* pipeline, which performs *minimap2* alignments individually for each read feature, and then haplotypes reads based on the combined annotations. With a *matchbox* script, this haplotyping can be performed in a single pass of the reads (Figure 6B). Applying the *matchbox* script to a healthy B-lymphoblastoid cell line HG02185 (20) as well as Cas9-targeted sequencing for FSHD1 sample 17706, we reproduced the haplotyping results of Xiao *et al*., identifying the pathogenic 4qA allele in 17706 as well as the presence of three different 4qA alleles indicating mosaicism (Figure 6C). Haplotyping of HG02185 also correctly identified the correct number of repeats and A/B haplotype of non-pathogenic alleles. In summary, for performing custom analysis or developing pipelines involving specific loci, *matchbox* scripts afford precision while remaining efficient and concise.

**Fig. 6.**
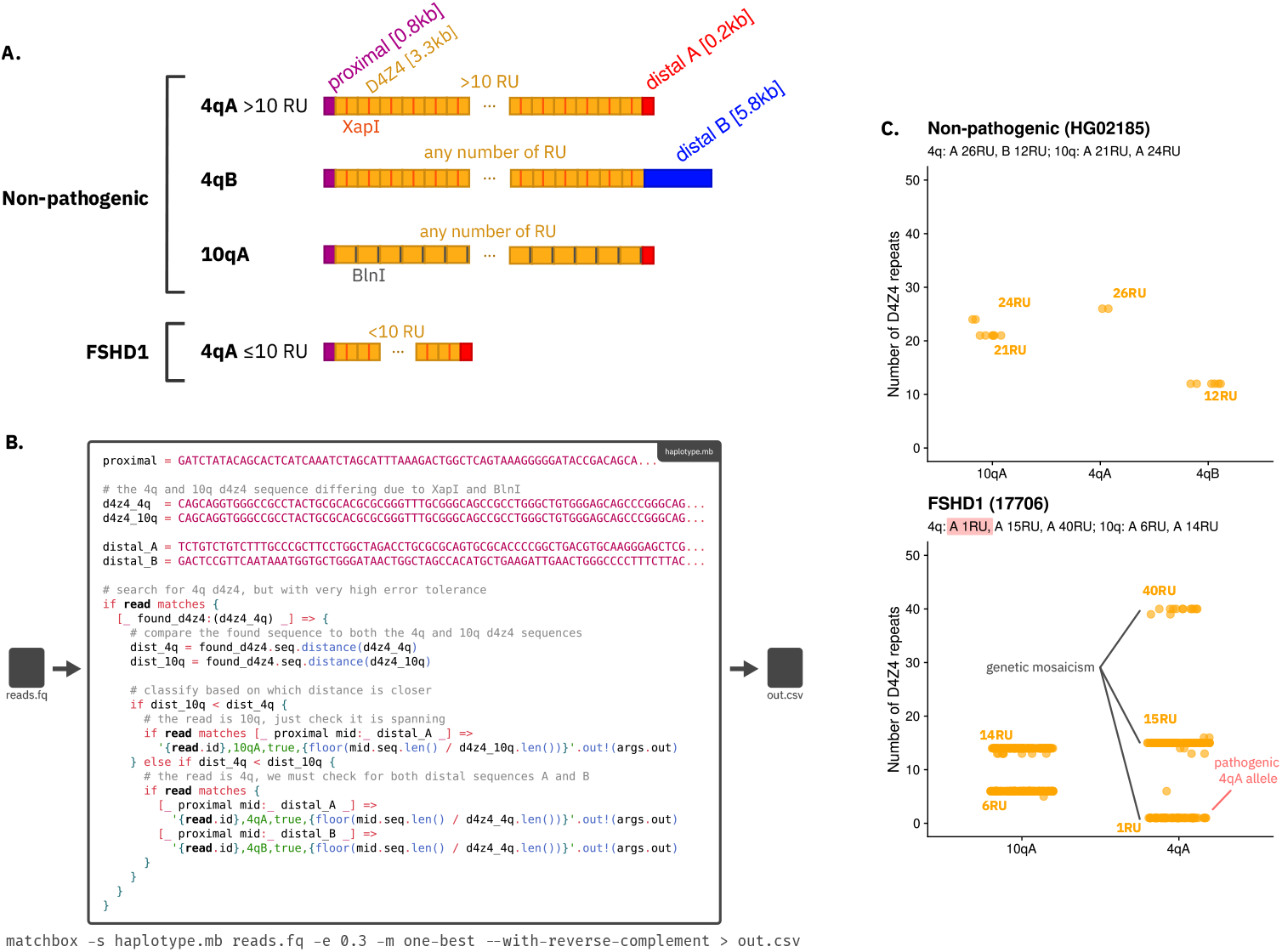
Application of *matchbox* to annotation and haplotyping of D4Z4 long-reads. (A) The structure of the D4Z4 array. The array contains a constant proximal region (purple), followed by a variable number of D4Z4 repeat units (RU, yellow) and a distal sequence (red or blue). The 4q array and homologous 10q array can be distinguished by differences in the sequence of the D4Z4 repeat, in particular the presence of XapI and BlnI restriction enzyme sites in the 4q and 10q alleles respectively. The 4qA and 4qB haplotypes can be distinguished by different sequences at the distal end of the array. (B) *matchbox* script for annotating and haplotyping D4Z4 reads. First, a D4Z4 repeat is located, and it is determined whether the sequence is closer to the 10q or 4q D4Z4. Then, the proximal and distal sequences are located, and the repeats are counted by dividing the area between the distal and proximal sequences by the length of the D4Z4 unit (3.3kb). An alternative method for counting repeats based on D4Z4 sequence matches was also attempted (Supplementary Fig. 2). The script outputs CSV results, recording each read’s ID, haplotype, and RUs. The parameter `-with-reverse-complement` allows every read to be processed along with its reverse-complement, allowing the user to only specify the pattern in the forward direction for concision. (C) The results of the script when applied to a healthy B-lymphoblastoid cell line HG02185 (top), and an FSHD patient sample 17706 (bottom).

### I. Computational performance

*matchbox*’s throughput varies significantly based on the script being run. Hence, a variety of benchmark tasks representing common read processing scenarios were devised to assess *matchbox*’s throughput, comparing against existing tools where appropriate. *matchbox* performed comparably to existing tools, in many cases outperforming purpose-built tools such as *Restrander* and *scTagger* (Figure 7). In particular, *matchbox* was the fastest tool for exact-match sequence searching (in this example, searching for a TSO primer sequence on either strand of a read). *matchbox*’s slowest task was the inference of primer arrangements in reads; this is likely due to the implementation of the `describe()` function forcing its computation to be performed at runtime, as opposed to *matchbox*’s pattern matching features which can preallocate their structures at compile time. Some established tools (*samtools, cutadapt, flexiplex, splitcode*) were able to outperform *matchbox* for some tasks, indicating that there are still optimisations to be made (Figure 7).

**Fig. 7.**
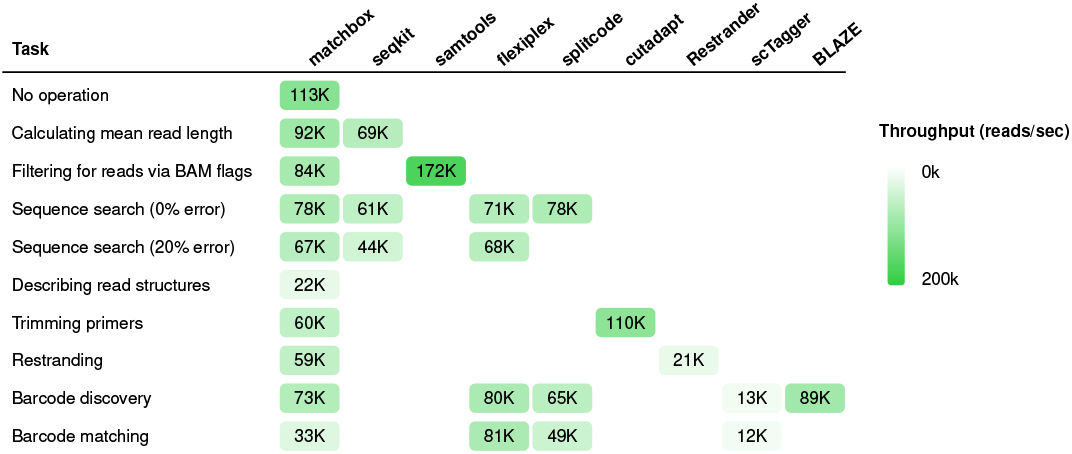
*matchbox* efficiently balances throughput (measured in reads/seq) and flexibility across a variety of read processing tasks. Each task was run 3 times on each tool and averaged. Samples of 10M reads were used.

## Discussion

*matchbox* is a powerful tool for navigating the complex read structures and fine-grained analysis needs of DNA and RNA sequencing data; support for protein sequence processing would involve only a minor modification and will be added in the future. *matchbox* is highly flexible in its potential applications, displays consistently fast computational performance over a wide variety of tasks and enables many complex analyses with just a few lines of code. The examples presented in this article are not a comprehensive account of *matchbox*’s applications, but an overview to highlight the tool’s general-purpose design.

*matchbox*’s syntax presents an ergonomic improvement over the limiting command-line parameters or configuration requirements of many existing tools, embracing the modularity that comes with a programming language to support user applications beyond what a tool’s author can conceive. However, a new textual programming language presents an inherent barrier, particularly to those with less experience on the command line. *matchbox*’s comprehensive documentation eases this problem, providing users with sample scripts for common applications and a full reference guide to *matchbox* features and functions. Future development of *matchbox* could also incorporate editor tooling such as syntax highlighting and static analysis to make the language more accessible, which would benefit both learners and experienced users. Another promising direction may be a visual interface for interactively devising preprocessing scripts. *matchbox*’s pattern-matching syntax, representing a read as a linear series of named regions, was already designed to more closely reflect the visual block-based depictions of read structures commonly deployed in the literature. Future development of an interactive *matchbox* interface could further unify the human readable diagrams used to represent reads with the pre-processing scripts used to investigate and manipulate them.

While *matchbox* has a suitably high throughput for large-scale data analysis, there remains room for further optimisations to be made in the tool’s implementation. The error-tolerant pattern-matching is the most expensive runtime operation and a more tailored implementation could improve performance, particularly for complex read structures with a lot of regions or barcodes to search for. Importantly, unlike for a task-specific program, any optimisations implemented within future updates to *matchbox* will benefit users across a wide variety of domains. *matchbox* can be installed via *cargo* with `cargo install matchbox-cli` .

## Author contributions statement

J.S. conceived of the project, developed the *matchbox* program, performed the analyses in this paper and wrote the manuscript. K.Z. assisted with the analyses and figure design, performed user testing and designed the *matchbox* logo. L.X. assisted with the D4Z4 analysis and figures. O.V. and S.J.V. facilitated the application of *matchbox* to CRISPR prime editing data, and S.M.R. generated datasets important for testing this application. M.E.R., Q.G. and M.B.C. planned and supervised the project and wrote the manuscript. All authors read and approved the final manuscript.

## ACKNOWLEDGEMENTS

We thank the WEHI Research Computing Platform for providing the Milton high performance computing platform. We thank Josh Zhang, Wendy Jia and Alex Yan for testing development versions of *matchbox* and providing feedback. We thank Sefi Prawer, Josie Gleeson, Andrew Wan, Yupei You, Changqing Wang, Callum Sargeant and Shian Su for helpful conversations during development which guided *matchbox* ‘s design. We also thank Brendan Zabarauskas for many helpful conversations and for his “language garden” repository, which strongly influenced the implementation of *matchbox*, especially its bidirectional type checking approach.

