## Supplementary methods, figures and tables for "Powerful read processing with *matchbox*"

### Powerful read processing with *matchbox*
#### Supplementary methods and figures

#### Contents


Supplementary methods ... 2
1. Basics ... 2
2. Demultiplexing ... 2
   2.1. 10X Discovery ... 2
   2.2. 10X Matching ... 2
   2.3. SPLiT-seq Matching ... 4
3. Restranding ... 4
   3.1. Correctness ... 4
   3.2. Complex *matchbox* script ... 5
   3.3. Restranding with trimming ... 8
   3.4. Protocols ... 9
4. CRISPR self-targeting screen ... 10
   4.1. Alternative approaches to determining edits ... 10
5. D4Z4 ... 11
   5.1. Length-based haplotyping ... 11

Supplementary figures ... 12
1. Editing efficiency in prime-editing screen data ... 12
2. Match-based D4Z4 haplotyping ... 12

Supplementary tables ... 13
1. Datasets ... 13
2. Software ... 13

Bibliography ... 14

### Supplementary methods

#### 1. Basics

To demonstrate *matchbox* on basic tasks, the PCB109 dataset was used. A random sample of 1M reads was created with *seqtk*.

```
seqtk sample ${reads} 1000000 > reads_1M.fq
```

#### 2. Demultiplexing

##### 2.1. 10X Discovery

For barcode discovery, the long-read compoment of scmixology2 was used. The GridION and PromethION components of HiPSC data generated by You *et al.* were also used, and true barcode lists for all three datasets came from You *et al.*'s *CellRanger* output. The full 10X barcode reference `3M-february-2018.txt` was provided to the demultiplexing tools, and the true barcode lists were used to check validity of discovered barcodes. In line with the parameter tuning in Cheng *et al.*, the number of expected cells for *BLAZE* was 200 for scmixology2 and 850 for the hiPSC datasets, and *flexiplex-filter* was given an upper search range of 1000 on hiPSC data.

```
# blaze
blaze ${reads} --threads=32 --expect-cells=${cells} --no-demultiplexing

# matchbox
matchbox -s discover_blaze.mb -a "out='discovered'" -o "." -e 0 -t 32 ${reads}
matchbox -s csv_to_tsv.mb -t 32 discovered.csv > discovered.tsv
flexiplex-filter -u 1000 -w 10x_barcode_set.csv -o filtered.tsv discovered.tsv

# scTagger
scTagger.py extract_lr_bc -t1 -r ${reads} -o "lr_output.tsv.gz"
pigz -dc lr_output.tsv.gz > lr_output.tsv
scTagger.py extract_sr_bc_from_lr -i "lr_output.tsv" -wl 3M-february-2018.txt -o
"discovered.tsv.gz"
pigz -dc sr_output.tsv.gz > sr_output.tsv
flexiplex-filter -u 1000 -w 3M-february-2018.txt -o filtered.tsv discovered.tsv

# flexiplex
flexiplex -f 0 -p 32 ${reads} > my_barcode_list.tsv
flexiplex-filter --whitelist 3M-february-2018.txt --outfile filtered.tsv
flexiplex_barcodes_counts.txt

# splitcode
echo "@extract {primer}<bc[16]>
group id  tag distance  maxFindsG
primers primer  CTACACGACGCTCTTCCGATCT  0 1" > splitcode.txt
splitcode -c splitcode.txt -t 32 ${reads} -o out.fq --empty-remove
matchbox -r "read.seq.count!(name='count')" bc.fastq -t 32
matchbox -r "'{read.value}\t{read.count}'.stdout!()" -t 32 count.csv > discovered.tsv
flexiplex-filter -u 1000 -w 3M-february-2018.txt -o filtered.tsv discovered.tsv
```

##### 2.2. 10X Matching

For barcode discovery, simulated data by Ebrahimi *et al.* was used, in particular the large simulated long-read component `L_lr.fa.gz`. The full list of 5000 true barcodes was extracted from the true-barcode-per-read TSV file `L.truth.tsv.gz`.

```
xan rename -n "id,seq" Ebrahimi_simulated/truth/L.truth.tsv.gz > ebrahimi_truth.csv
xan select seq ebrahimi_truth.csv | xan dedup > ebrahimi_full_set.csv
```

A subset of 500 barcodes was taken, and the reads corresponding to these 500 barcodes were subsampled.

```
xan sample 500 ebrahimi_full_set.csv > ebrahimi_500_set.csv
xan join seq ebrahimi_truth.csv seq ebrahimi_500_set.csv > ebrahimi_500_ids_seqs.csv
xan select id ebrahimi_500_ids_seqs.csv > ebrahimi_500_ids.csv
seqtk subseq Ebrahimi_simulated/long-reads/L_lr.fa.gz ebrahimi_500_ids.csv >
ebrahimi_subset.fa
```

*matchbox* was used to annotate each of these subsampled reads with its barcode:

```
# convert the CSV to a FASTA
matchbox --run "'>{read.id}\n{read.seq}'.stdout!()" ebrahimi_500_ids_seqs.csv >
ebrahimi_subset_barcodes.fa

# then run in paired mode
matchbox ebrahimi_subset.fa --paired-with ebrahimi_subset_barcodes.fa --run
"read.r1.tag('barcode={read.r2.seq}').out!('ebrahimi_subset_annotated.fa')"
```

Then each tool was run on the annotated file. In line with the analysis in Cheng *et al.*, ambiguous matches were removed from the *scTagger* output.

```
# flexiplex
flexiplex -d 10x3v3 -e ${editDistance} -k ebrahimi_full_set.csv -p 32 -n
flexiplex_${editDistance}_ ebrahimi_subset_annotated.fa > demultiplexed.fa
matchbox -s compare_flexiplex.mb -t 32 -a "out='flexiplex_${editDistance}'" demultiplexed.fa -
p ebrahimi_subset_annotated.fa

# matchbox
matchbox -s demultiplex.mb -t 32 -a "barcodes='ebrahimi_full_set.csv',
out='demultiplexed_${errorRate}.fa'" -e ${errorRate} -m one-best ebrahimi_subset_annotated.fa
matchbox -s compare_matchbox.mb -t 32 -a "out='matchbox_${errorRate}'" -e ${errorRate} -m one-
best demultiplexed_${errorRate}.fa

# scTagger
# first convert to FASTQ with blank qscore, scTagger will miss half the reads
matchbox --run "{ id=read.id, desc=read.desc, seq=read.seq, qual='' }.out!
('ebrahimi_subset_annotated.fq')"
# behead CSV because scTagger isn't expecting header
xan behead ebrahimi_full_set.csv > ebrahimi_full_set_headless.csv
scTagger.py extract_lr_bc -r redas.fq -t 32 -o temp.out.gz
pigz -d temp.out.gz
scTagger.py match_trie -mr ${editDistance} -lr temp.out -sr barcodes_headless.csv -t 32 -o
out.txt
# filter out ambiguous matches, as in Cheng et al
xan rename id,edit_distance,matches,segment,barcode out.txt -d "\t" | xan filter "matches==1"
| xan select id,barcode > out.csv
xan join id out.csv id ebrahimi_500_ids_seqs.csv > joined.csv
matchbox -s compare_sctagger.mb -t 32 -a "out='sctagger_${editDistance}'" joined.csv

# splitcode
# (can't get splitcode to work well on reverse reads,
# so let matchbox help out by restranding first)
matchbox -m one-best -e 0.2 -r "primer = CTACACGACGCTCTTCCGATCT; if read matches {[_ primer _]
=> read.out!('restranded.fa'), [_ -primer _] => (-read).out!('restranded.fa')}" ${readsFaGz}
echo "group   id tag  distance    minFindsG   maxFindsG   next
prims   primer  CTACACGACGCTCTTCCGATCT  3:3:3    1   1   {barcode}0-0
bc  barcode barcodes.txt    ${editDistance}:${editDistance}:${editDistance}   1   1" >
splitcode.txt
# make the forward and reverse barcode files
cp ${barcodesCsv} barcodes.txt
# run splitcode
splitcode -c splitcode.txt -t 32 -o out.fq --mod-names --seq-names restranded.fa
```

*matchbox* scripts were written to check the count the correct barcodes assigned by each tool against the reference.

##### 2.3. SPLiT-seq Matching

The long-read SPLiT-seq dataset generated by Rebboah *et al.* was used. The TSV outputs were joined and further analysis was performed in R to compare the barcodes, checking for an exact match in each of the three barcode regions.

```
# LR-splitpipe
python3 demul all -f ${reads} -o splitpipe -k WT -c v2 -t 32 --max_linker_dist 1000 --
chunksize 10000

# matchbox
# (altered script to output TSVs)
matchbox -s demultiplex.mb -e 0.2 ${reads} -t 32 -m one-best > bcs_headless.tsv
xan rename read_name,mb_bc1,mb_bc2,mb_bc3,mb_umi bcs_headless.tsv -n > matchbox_out.tsv

# join the outputs
xan join read_name matchbox_out.tsv read_name splitpipe_bcs.tsv > joined.tsv
```

#### 3. Restranding

##### 3.1. Correctness

Dong *et al.* PCB109 dataset was used, including spike-in sequins.

For panel C, reads were aligned to a reference file `rnasequin_sequences_2.4.fa` containing the sequences for the RNA Sequins.

```
minimap2 -ax splice:hq -o aligned.sam -t 32 rnasequin_sequences_2.4.fa reads.fq.gz
```

Then, a *matchbox* script was used to parse the SAM file, annotating each aligned read with its strand and outputting in FASTQ format.

```
# filter for aligned reads
if read.seq != '*' and read.rname != '*' {
  # create a copy of the read with an empty description field
  new_read = {
    id = read.id,
    seq = read.seq,
    qual = read.qual,
    desc = ''
  }

  # then, tag it based on minimap2 alignment
  if read.flag.reverse_strand() {
    new_read.tag('true_strand=-').out!(args.out)
  } else {
    new_read.tag('true_strand=+').out!(args.out)
  }
}
```

```
matchbox -s annotate.mb -t 32 -a "out = 'annotated.fq'" aligned.sam
```

Then `annotated.fq` was restranded with the various restranding methods.

For *matchbox* and *Restrander*, an error rate of 0.25 was used for primer sequences. *pychopper* only provides an auto mode, which determines the best error rate automatically. The *pychopper* `edlib` backend was used, as it has been previously shown to be more effective and faster than the machine learning backend [1].

```
pychopper -m edlib

restrander annotated.fq restranded.fq PCB109.json

matchbox -s restrand_simple.mb -e 0.25 annotated.fq -t 32 -m one-best -a "
```

```
  tso = TTTCTGTTGGTGCTGATATTGCTGGG,
  rtp = ACTTGCCTGTCGCTCTATCTTCTTTTTTTTTT,
  out = 'restranded.fq',
  unknowns_out = 'unknowns.fq',
  name = '0.25'
"

matchbox -s restrand_complex.mb -e 0.25 annotated.fq -t 32 -m one-best -a "
  tso = TTTCTGTTGGTGCTGATATTGCTGGG,
  rtp = ACTTGCCTGTCGCTCTATCTTCTTTTTTTTTT,
  out = 'restranded.fq',
  unknowns_out = 'unknowns.fq',
  name = '0.25'
"
```

##### 3.2. Complex *matchbox* script

The `matchbox_complex.mb` script is more verbose than the basic script, closely following the program logic of *Restrander* to almost perfectly replicate its classifications.

```
tso = args.tso
rtp = args.rtp

polyA = AAAAAAAAAAAA

count!('total', name=args.name)

if read matches {
  [_ |200:(_ polyA~0 _)|] => {
    if read matches {
      [|200:(_ (-polyA)~0 _)| _] => {
        if read matches {
          [|200:(_ tso _)| _] => {
            if read matches {
              [_ -tso _] => {
                count!('tso-tso artefact', name=args.name)
                read.tag('strand=?').out!(args.unknowns_out)
              }
              [|200:(_ rtp _)| _] => {
                count!('?', name=args.name)
                read.tag('strand=?').out!(args.unknowns_out)
              }
              [_] => {
                count!('+', name=args.name)
                read.tag('strand=+').out!(args.out)
              }
            }
          }
          [|200:(_ rtp _)| _] => {
            if read matches {
              [_ -rtp _] => {
                count!('rtp-rtp artefact', name=args.name)
                read.tag('strand=?').out!(args.unknowns_out)
              }
              [|200:(_ tso _)| _] => {
                count!('?', name=args.name)
                read.tag('strand=?').out!(args.unknowns_out)
              }
              [_] => {
                count!('-', name=args.name)
                read.tag('strand=-').out!(args.out)
              }
            }
          }
          [_] => {
            count!('?', name=args.name)
            read.tag('strand=?').out!(args.unknowns_out)
          }
        }
      }
      [_] => {
        count!('+', name=args.name)
        read.tag('strand=+').out!(args.out)
      }
    }
```

```
}
[|200:(_ (-polyA)~0 _)| _] => {
  if read matches {
    [_ |200:(_ polyA~0 _)|] => {
      if read matches {
        [|200:(_ tso _)| _] => {
          if read matches {
            [_ -tso _] => {
              count!('tso-tso artefact', name=args.name)
              read.tag('strand=?').out!(args.unknowns_out)
            }
            [|200:(_ rtp _)| _] => {
              count!('?', name=args.name)
              read.tag('strand=?').out!(args.unknowns_out)
            }
            [_] => {
              count!('+', name=args.name)
              read.tag('strand=+').out!(args.out)
            }
          }
        }
        [|200:(_ rtp _)| _] => {
          if read matches {
            [_ -rtp _] => {
              count!('rtp-rtp artefact', name=args.name)
              read.tag('strand=?').out!(args.unknowns_out)
            }
            [|200:(_ tso _)| _] => {
              count!('?', name=args.name)
              read.tag('strand=?').out!(args.unknowns_out)
            }
            [_] => {
              count!('-', name=args.name)
              read.tag('strand=-').out!(args.out)
            }
          }
        }
        [_] => {
          count!('?', name=args.name)
          read.tag('strand=?').out!(args.unknowns_out)
        }
      }
    }
    [_] => {
      count!('-', name=args.name)
      read.tag('strand=-').out!(args.out)
    }
  }
}
[_] => {
  if read matches {
    [|200:(_ tso _)| _] => {
      if read matches {
        [_ -tso _] => {
          count!('tso-tso artefact', name=args.name)
```

```
            read.tag('strand=?').out!(args.unknowns_out)
          }
          [|200:(_ rtp _)| _] => {
            count!('?', name=args.name)
            read.tag('strand=?').out!(args.unknowns_out)
          }
          [_] => {
            count!('+', name=args.name)
            read.tag('strand=+').out!(args.out)
          }
        }
      }
      [|200:(_ rtp _)| _] => {
        if read matches {
          [_ -rtp _] => {
            count!('rtp-rtp artefact', name=args.name)
            read.tag('strand=?').out!(args.unknowns_out)
          }
          [|200:(_ tso _)| _] => {
            count!('?', name=args.name)
            read.tag('strand=?').out!(args.unknowns_out)
          }
          [_] => {
            count!('-', name=args.name)
            read.tag('strand=-').out!(args.out)
          }
        }
      }
      [_] => {
        count!('?', name=args.name)
        read.tag('strand=?').out!(args.unknowns_out)
      }
    }
  }
}
```

Then, a *matchbox* script was applied to count the number of correct and incorrect restranding classifications by comparing the tags in each read's description.

```
# locate the 'true_strand' tag
a = read.desc.find_first('true_strand=')
ts = read.desc.slice(a + 13, a + 14)

# locate the 'strand' tag
b = read.desc.find_first(' strand=')
ps = read.desc.slice(b + 8, b + 9)

# compare the two
(ts == ps).count!(name=args.name)
```

3.3. Restranding with trimming

The script shown in 4B can be modified to additionally trim reads:

```
# define the primer sequences
tso = TTTCTGTTGGTGCTGATATTGCTGGG
rtp = ACTTGCCTGTCGCTCTATCTTCTTTTTTTTTT

if read matches {
  # first, check if read is an amplification artefact
  [_ tso _ -tso _] =>
    { read.tag('strand=?').out!(args.unknown); count!('tso artefact') }
  [_ rtp _ -rtp _] =>
    { read.tag('strand=?').out!(args.unknown); count!('rtp artefact') }

  # then, separate forward and reverse reads
  [_ tso trimmed:_ -rtp _] =>
    { trimmed.tag('strand=+').out!(args.out); count!('+') }
  [_ rtp trimmed:_ -tso _] =>
    { (-trimmed).tag('strand=-').out!(args.out); count!('-') }

  # keep uncategorised reads, to investigate them further
  [_] => { read.tag('strand=?').out!(args.unknown); count!('?') }
}
```

##### 3.4. Protocols

The *matchbox* script was applied to a number of protocols. Datasets used were Tian *et al.* (MSCv2) [2], Dong *et al.* (PCB109) [3], Subas Satish *et al.* (NEBNext) [4], and Schuster *et al.* (nanoslam) [1].

```
# MSCv2
restrander MSCv2.fq MSCv2_restranded.fq config/10X-3prime.json > stats.json
matchbox -s restrand_complex.mb -e 0.25 MSCv2.fq -t 32 -m one-best -a "
  tso = TTTCTGTTGGTGCTGATATTGCTGGG,
  rtp = ACTTGCCTGTCGCTCTATCTTCTTTTTTTTTT,
  out = 'MSCv2_restranded.fq',
  unknowns_out = 'MSCv2_restranded.fq',
  name = 'MSCv2'
"

# NEBNext
restrander NEBNext.fq NEBNext_restranded.fq config/NEBNext.json
matchbox -s restrand_complex.mb -e 0.25 NEBNext.fq -t 32 -m one-best -a "
  tso = GCTAATCATTGCAAGCAGTGGTATCAACGCAGAGTACAT,
  rtp = AAGCAGTGGTATCAACGCAGAGTACTTTTTTTTTTTTTTTTTTTTTTTTTTTTTT,
  out = 'NEBNext_restranded.fq',
  unknowns_out = 'NEBNext_restranded.fq',
  name = 'NEBNext'
"

# nanoslam
restrander nanoslam.fq nanoslam_restranded.fq config/PCB111.json
matchbox -s restrand_complex.mb -e 0.25 NEBNext.fq -t 32 -m one-best -a "
  tso = TTTCTGTTGGTGCTGATATTGCTTT,
  rtp = CTTGCCTGTCGCTCTATCTTCAGAGGAG,
  out = 'nanoslam_restranded.fq',
  unknowns_out = 'nanoslam_restranded.fq',
  name = 'nanoslam'
"

# PCB109
restrander PCB109.fq PCB109_restranded.fq config/PCB109.json
matchbox -s restrand_complex.mb -e 0.25 NEBNext.fq -t 32 -m one-best -a "
  tso = TTTCTGTTGGTGCTGATATTGCTGGG,
  rtp = ACTTGCCTGTCGCTCTATCTTCTTTTTTTTTT,
  out = 'PCB109_restranded.fq',
```

```
  unknowns_out = 'PCB109_restranded.fq',
  name = 'PCB109'
"
```

#### 4. CRISPR self-targeting screen

Reads from Cirincione and Simpson *et al.* were downloaded corresponding to both cell lines, all time points and both replicates of the self-targeting 'sensor' libraries. Supplementary Table 1 of Cirincione and Simpson *et al.* was retrieved from the spreadsheet `41592_2024_2502_MOESM3_ESM.xlsx`, converted to a CSV file `refs.csv` and used as a barcode reference file.

| Accession | Cell line | Day | Replicate |
|---|---|---|---|
| SRR30675759 | PEmax | 7 | 1 |
| SRR30675758 | PEmax | 7 | 2 |
| SRR30675749 | PEmax | 14 | 1 |
| SRR30675748 | PEmax | 14 | 2 |
| SRR30675745 | PEmax | 21 | 1 |
| SRR30675744 | PEmax | 21 | 2 |
| SRR30675755 | PEmax | 28 | 1 |
| SRR30675754 | PEmax | 28 | 2 |
| SRR30675751 | PEmaxKO | 7 | 1 |
| SRR30675750 | PEmaxKO | 7 | 2 |
| SRR30675747 | PEmaxKO | 14 | 1 |
| SRR30675746 | PEmaxKO | 14 | 2 |
| SRR30675757 | PEmaxKO | 21 | 1 |
| SRR30675756 | PEmaxKO | 21 | 2 |
| SRR30675753 | PEmaxKO | 28 | 1 |
| SRR30675752 | PEmaxKO | 28 | 2 |

The `quantify_edits.mb` script was run on each sample, and all further filtering and visualisation was performed in *R*.

```
matchbox ${cell_line}_${day}_${replicate}.fq  -s quantify_edits.mb r2.fq -o "${cell_line}
_${day}_${replicate}"
```

Figures D and E are condensed recapitulations of Fig 2c and 2d from Cirincione and Simpson *et al.* A more direct reproduction is included here.

##### 4.1. Alternative approaches to determining edits

When determining whether a read has been edited, one method alternative to 5C would be to discard all reads with off-target edits, only counting reads which exactly match either the unedited or edited sequence.

```
refs = csv(args.refs)

if read matches [_ target:|47| -ref.barcode AAAAAAA _] for ref in refs => {
  # check if the sequence is edited or unedited
  if target.seq == ref.unedited {
    average!(0, name=ref.barcode)
  } else if target.seq == ref.edited {
    average!(1, name=ref.barcode)
  }
}
```

Another approach would simply be to quantify occurrences of each exact sequence detected; although random sequencing error will lead to a large number of different sequences, if a guide is consistently performing an edit, this will show up.

```
refs = csv(args.refs)

if read matches [_ target:|47| -ref.barcode AAAAAAA _] for ref in refs => {
  # simply count the found sequences
  count!(target.seq, name=ref.barcode)
}
```

## 5. D4Z4

##### 5.1. Length-based haplotyping

It should be noted that there is a fixed sequence between the proximal sequence and the first D4Z4 repeat (1kb) and an additional partial D4Z4 repeat at the distal end of the array (of varying length, up to 1.9kb). Because of this, the calculation `mid.seq.len() / d4z4_4q.len()` (or `mid.seq.len() / d4z4_10q.len()`) is always an overestimation of RU, by at most 2.9kb (0.87 RU). Hence, flooring the result with the `floor()` function (as opposed to rounding it with `round()`) provides the best estimate of RU.

### Supplementary figures

1. Editing efficiency in prime-editing screen data

2. Match-based D4Z4 haplotyping

A.

reads.fq → haplotype_match.mb → out.csv

```
proximal = GATCTATACAGCACTCATCAAATCTAGCATTTAAAGACTGGCTCAGTAAAGGGGGATACCGAC...

# the 4q and 10q d4z4 sequence differing due to XapI and BlnI
d4z4_4q  = CAGCAGGTGGGCCGCCTACTGCGCACGCGGGTTTGCGGGCAGCCGCCTGGGCTGTGGGAGCAGCCCGGGCAG...
d4z4_10q = CAGCAGGTGGGCCGCCTACTGCGCACGCGGGTTTGCGGGCAGCCGCCTGGGCTGTGGGAGCAGCCCGGGCAG...

distal_A = TCTGTCTGTCTTTGCCCGCTTCCTGGCTAGACCTGCGCGCAGTGCGCACCCCGGCTGACGTGCAAGGGAGCTCG...
distal_B = GACTCCGTTCAATAAATGGTGCTGGGATAACTGGCTAGCCACATGCTGAAGATTGAACTGGGCCCCTTTCTTAC...

if read.qual.to_qscores().mean() > 14 {
  # search for 4q d4z4, but with very high error  tolerance
  if read matches {
    [_ found_d4z4:(d4z4_4q) _] => {
      dist_4q = found_d4z4.seq.distance(d4z4_4q)
      dist_10q = found_d4z4.seq.distance(d4z4_10q)

      if dist_10q < dist_4q {
        if read matches [_ p13E11 mid:_ distal_A _] =>
          stdout!('{read.id},10qA,true,{mid.count_matches(d4z4_10q, error_rate=0.2)}')
      } else if dist_4q < dist_10q {
        if read matches {
          [_ p13E11 mid:_ distal_A _] =>
            stdout!('{read.id},4qA,true,{mid.count_matches(d4z4_4q, error_rate=0.2)}')
          [_ p13E11 mid:_ distal_B _] =>
            stdout!('{read.id},4qB,true,{mid.count_matches(d4z4_4q, error_rate=0.2)}')
        }
      }
    }
  }
}
```

```
matchbox -s haplotype.mb reads.fq -e 0.3 -m one-best --with-reverse-complement > out.csv
```

B.

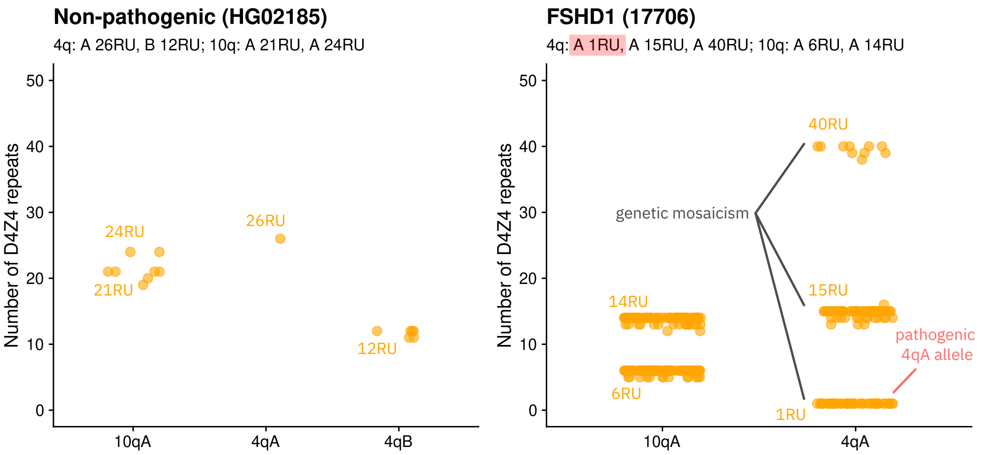


Figure 1: Application of *matchbox* to annotation and haplotyping of D4Z4 reads, modified to count the number of D4Z4 sequences found, rather than estimate it using the length of the region between the distal sequences. (A) The *matchbox* script for haplotyping. Instead of approximating repeats based on length, the `count_matches()` function is used. (B) The results of the script when applied to a healthy B-lymphoblastoid cell line HG02185 (left), and an FSHD patient sample 17706 (right).

### Supplementary tables

#### 1. Datasets

| Accession | Citation | Platform | Resolution | Used for |
|---|---|---|---|---|
| GEO GSE172421 | Dong *et al.* [3] | ONT | Bulk | Sequence search, restranding |
| GEO GSE154870 | Tian *et al.* [2] | ONT | 10X single-cell | Barcode discovery, restranding |
| ENA PRJEB54718 | You *et al.* [5] | ONT | 10X single-cell | Barcode discovery |
| 10.6084/ m9.figshare.19740475.v1 | Ebrahimi *et al.* [6] | Simulated long-reads | 10X single-cell | Barcode matching |
| SRR13948564 | Rebboah *et al.* [7] | PacBio | SPLiT-seq single-cell | Barcode matching |
| ENA PRJEB51442 | Subas Satish *et al.* [4] | ONT | Bulk | Restranding |
| ENA PRJEB60282 | Schuster *et al.* [1] | ONT | Bulk | Restranding |
| ENA PRJNA1159206 | Cirincione and Simpson *et al.* [8] | Illumina | Bulk | CRISPR screen analysis |
| EGAD50000001551 | Xiao *et al.* [9] | ONT | Bulk | D4Z4 haplotyping |
| 1KGP-ONT HG02185 | Gustafson *et al.* [10] | ONT | Bulk | D4Z4 haplotyping |

Table 1: List of datasets used to demonstrate *matchbox*.

#### 2. Software

| Software | Task | Details |
|---|---|---|
| *Rust-Bio* v3.0.0 [11] | Implementation | *matchbox* sequence search algorithms |
| *rayon* v1.11.0 [12] | Implementation | *matchbox* multi-threaded execution |
| *flexiplex* v1.02.4 [13] | Benchmarking | Sequence search, discovery and matching of 10X barcodes |
| *cutadapt* v4.9 [14] | Benchmarking | Sequence search, primer trimming |
| *splitcode* v0.31.2 [15] | Benchmarking | Sequence search, discovery and matching of 10X barcodes |
| *scTagger* v1.1.1 [6] | Benchmarking | Discovery and matching of 10X barcodes |
| *BLAZE* v2.4.0 [5] | Benchmarking | Discovery of 10X barcodes |
| *LR-splitpipe* v2.0 [7] | Benchmarking | Demultiplexing of SPLiT-seq barcodes |
| *Restrander* v1.1.1 [1] | Benchmarking | Restranding PCR-cDNA reads |
| *pychopper* v2.7.10 [16] | Benchmarking | Restranding PCR-cDNA reads |
| *D4Z4End2End* [9] | Benchmarking | Annotating and haplotyping raw D4Z4 reads |
| *samtools* v1.22.1 [17] | Benchmarking, analysis | General SAM/BAM processing |
| *seqtk* v1.5 [18] | Benchmarking, analysis | Sequence search |
| *seqkit* v2.10.1 [19] | Benchmarking | Sequence search |
| *conda* v24.7.1 [20] | Analysis | Environment management |
| *Nextflow* v23.10.0 [21] | Analysis | Workflow management |
| *hyperfine* v1.19.0 [22] | Analysis | Profiling |
| *R* v4.5.1 [23] | Analysis | Data analysis and visualisation |
| *dplyr* v1.1.4 [24] | Analysis | Data analysis |
| *ggplot2* v3.5.2 [25] | Analysis | Data visualisation |

Table 2: List of software tools most integral to creating and demonstrating *matchbox*. "Implementation" refers to software used internally in the implementation of *matchbox*. "Benchmarking" refers to software *matchbox* was benchmarked against. "Analysis" and "visualisation" refer to software used in the processing and presentation of analysis throughout the paper.

[22] D. Peter, “hyperfine.” [Online]. Available: https://github.com/sharkdp/hyperfine

[23] R Core Team, *R: A Language and Environment for Statistical Computing*. Vienna, Austria: R Foundation for Statistical Computing, 2018. [Online]. Available: https://www.r-project.org/

[24] H. Wickham, R. François, L. Henry, K. Müller, and D. Vaughan, *dplyr: A Grammar of Data Manipulation*. 2025. [Online]. Available: https://dplyr.tidyverse.org/

[25] H. Wickham, *Ggplot2: Elegant graphics for data analysis*, 2nd ed. in Use R!. Cham, Switzerland: Springer International Publishing, 2016.
